# Skewed distributions of LRS: beyond mean and variance

**DOI:** 10.1101/696500

**Authors:** Shripad Tuljapurkar, Wenyun Zuo, Tim Coulson, Carol Horvitz, Jean-Michel Gaillard

## Abstract

Many field studies find that lifetime reproductive success (LRS) is highly skewed and often multimodal among individuals. Field biologists generate invaluable data on survival and reproductive rates, as a function of age and stage, that are used to parameterise structured models. These models often perform well at predicting population growth and mean LRS, but we do not know whether they accurately predict observed distributions of individual LRS. If the models fail to recreate these distributions, their use may be limited because the LRS is central to understanding life history evolution. We present powerful tools to generate distributions of LRS from age and/or stage structured models. Our methods reveal that structured models do perform well at generating distributions that agree with observations. Our approach also reveals why such skewed distributions arise, and helps resolve a debate about detecting signatures of selection in skewed distributions of LRS.

## Introduction

Lifetime measures of reproductive performance describe the number of off-spring an individual produces over its lifespan (Clutton-Brock 1988). Many empirical biologists consider such lifetime measures as individual fitness (e.g. Kruuk et al. 1999; Brommer et al. 2004), although, with the exception of annual species (Fisher 1930), these measures are correlates of fitness (Grafen 1988). Nonetheless, lifetime reproductive performance can provide generational estimators of the opportunity for natural and sexual selection and phenotypic evolution. For instance, many empirical studies have assessed the opportunity for selection using the standardized variance (I metric, Crow 1958) of reproductive performance and revealed high variation (e.g. in females from 0.177 in lions, *Panthera leo* to 3.62 in elephant seals, *Mirounga angustirostris* (Cabana and Kramer 1991); and in males from 0.59 in Atlantic salmon *Salmo salar* to 4.52 in bighorn sheep, *Ovis canadensis*, (Tatarenkov et al. 2008)). Comparison of standardized variances of lifetime reproductive performance between sexes within a population is commonly used for assessing the strength of sexual selection (e.g. Dubuc et al. (2014) on Rhesus macaques, *Macaca mulatta*). These empirical studies consistently reveal that distributions of lifetime reproductive performance measures are often non-normal, zero-inflated and highly skewed, and this begs the question: what are the demographic causes of these unusually-shaped distributions?

Lifetime Breeding Success (LBS) censuses offspring at birth (e.g. Slate et al. 2000), while Lifetime Reproductive Success (LRS) censuses them at some later stage (e.g. weaning/fledging (van Noordwijk and van Schaik 1999), one year-old (Newton 1988), two years-old (Clutton-Brock 1988)). We analyze discrete-time population models that are formulated in terms of species-appropriate modes of breeding and counting, but for simplicity use just the term LRS. So the question is: do such models predict LRS distributions that mimic those observed in nature, and why?

In the models here, individuals are distinguished by age and/or stage, and life cycles are described by probabilities of survival, stage transitions, and reproduction. Simulations for many species yield a distribution of LRS that is dramatically skewed or multimodal (Tuljapurkar et al. 2009b; Steiner and Tuljapurkar 2012), in accord with the empirical results mentioned above. Hence in any cohort, many individuals have an LRS of 0 so they leave no offspring; a lucky few have an LRS much higher than the average value. This variation is driven purely by chance since all the individuals are governed by the same probability rules (Steiner and Tuljapurkar 2012; van Daalen and Caswell 2017) (hereafter the latter reference is abbreviated vDC), meaning that “luck” can be more important than “pluck” (Snyder and Ellner 2018).

To quantify such “neutral” variation, Steiner and Tuljapurkar (2012) obtained formulas for the mean and variance (and higher moments) using a fixed fertility at every age/stage. But, as emphasized by vDC, offspring only come in integers (0, 1, 2, …), so reproduction at every age/stage should properly be described by probabilities that an individual has 0, 1, 2, …off-spring. Using the latter description, vDC derived formulas for the moments mean, variance, and so on – of LRS for each birth stage (extending Caswell (2011) and van Daalen and Caswell (2015)). The analyses by Snyder and Ellner (2018) also examined variance computed in this way as a measure of dispersion in the LRS. For populations structured only by age, the variance in LRS has previously been computed by Waples et al. (2011) and Charlesworth and Williamson (1975).

Yet, the moments do not provide the same information as the probability distribution of the LRS. Why? After all, the formulas in vDC easily yield the first three moments (also higher order moments but these don’t reveal such interesting patterns as multimodality). For one thing, our intuitive understanding of these moments is often based on a roughly normal distribution and so can be of limited value. For another, a general approach to approximating a distribution by the first few moments, e.g., the Gram-Charlier expansion (Stuart et al. 1994), doesn’t work for many (all the ones we’ve tried) LRS distributions, as we illustrate here. Thus the complete set of probabilities for LRS is often essential, certainly when the distribution is highly skewed or multimodal.

We present here a new method for computing the entire probability distribution of LRS for most species. We show below by examples that the distribution of LRS contains new and useful biological insights. Our methods highlight important questions: what factors determine the probability that individuals produce zero offspring during their lifetimes; how does a skewed LRS distribution affect natural selection, effective population size, and life history evolution? We discuss these questions, and point towards future work needed in this area.

The main text explains our method, and then focuses on examples. The first two examples are age-only models for human populations: the Hadza, a Tanzanian group of hunter-foragers, and a North American group called the Hutterites. The next example is a stage-only model for the large evergreen tree *Tsuga canadensis*. For these examples, the sources are cited in vDC, who also kindly provided matrices. The last example, and perhaps most general, is an age+stage model for Roe deer, *Capreolus capreolus*, based on the data in Plard et al. (2015). For each example we compute and display the full distribution, and also the first 3 moments (the same as in vDC).

## Methods

Lifetime Reproductive Success (LRS) is a random quantity, and we find the probability for each possible value of LRS. We model demography in discrete time; vital rates may be structured only by age, only by stage, or by a combination of age and stage (age+stage). Mathematical details are in the online Appendix; we use and explain discrete convolutions and discrete Fourier transforms, unfamiliar to some biologists, but essential here. Code (based on R (R Core Team 2019) and the R package FFTW (Fastest Fourier Transform in the West) (Mersmann 2019)) is available from the authors. Our results should apply to continuous stages using established methods (Ellner et al. 2016; Snyder and Ellner 2018). Notation is always a nuisance; where possible, we use notation similar to Caswell (2011) and Caswell et al. (2018).

Our logic uses three points. First, every individual has a particular age of death; second, at each age before death, there is a probability distribution for the number of offspring produced at that age; and third, convolution of these age-specific distributions (weighted appropriately by mortality) yields the distribution of LRS. We make the assumption that the probability distribution at each age is independent of the distribution at all other ages.

### General Framework

Age is counted in discrete intervals *a* = 1, 2, …, *ω* and newborns are always in the first age interval (parameters are defined in Table 1). Stage is defined in discrete categories, *s* = 1, 2, …, *S* (indexed as *i* or *j*) for stage-only and age+stage models. Stage transition probabilities (conditional on remaining alive) over a single time step are defined in a matrix ***G*** for stage-only models, or in a set of matrices ***G***_*a*_ for age+stage models. One-period survival probabilities (from one interval of time to the next) are *p*_*a*_ for age-only models, *p*(*j*) for stage-only models and *p*_*a*_(*j*) for age+stage models (Table 1). Multiplying stage transitions by stage survivals, we obtain an *S* × *S* matrix of unconditional stage transition probabilities, ***U*** for stage-only models and a set ***U***_*a*_ for age+stage models (Table 1).

**Table 1:**
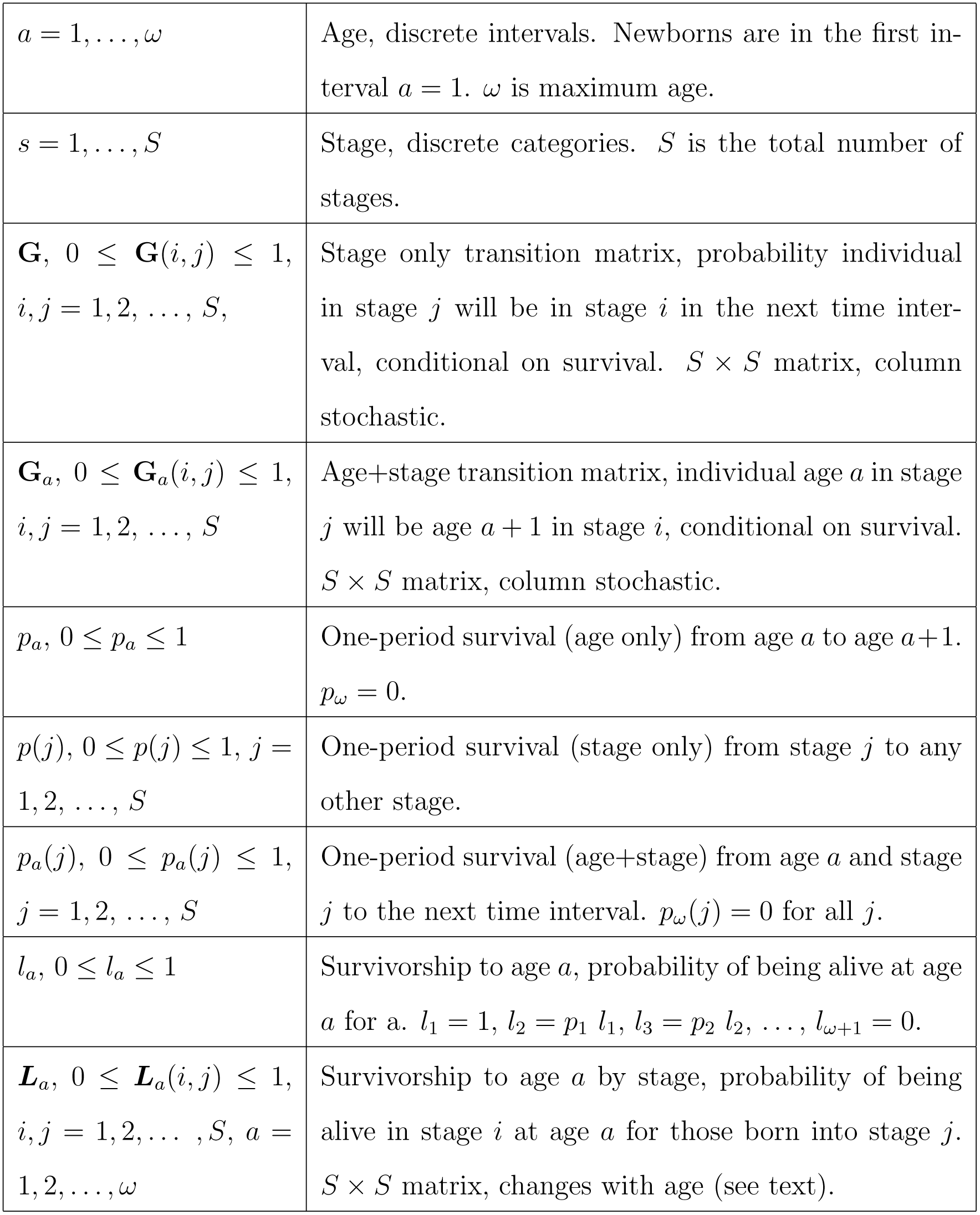
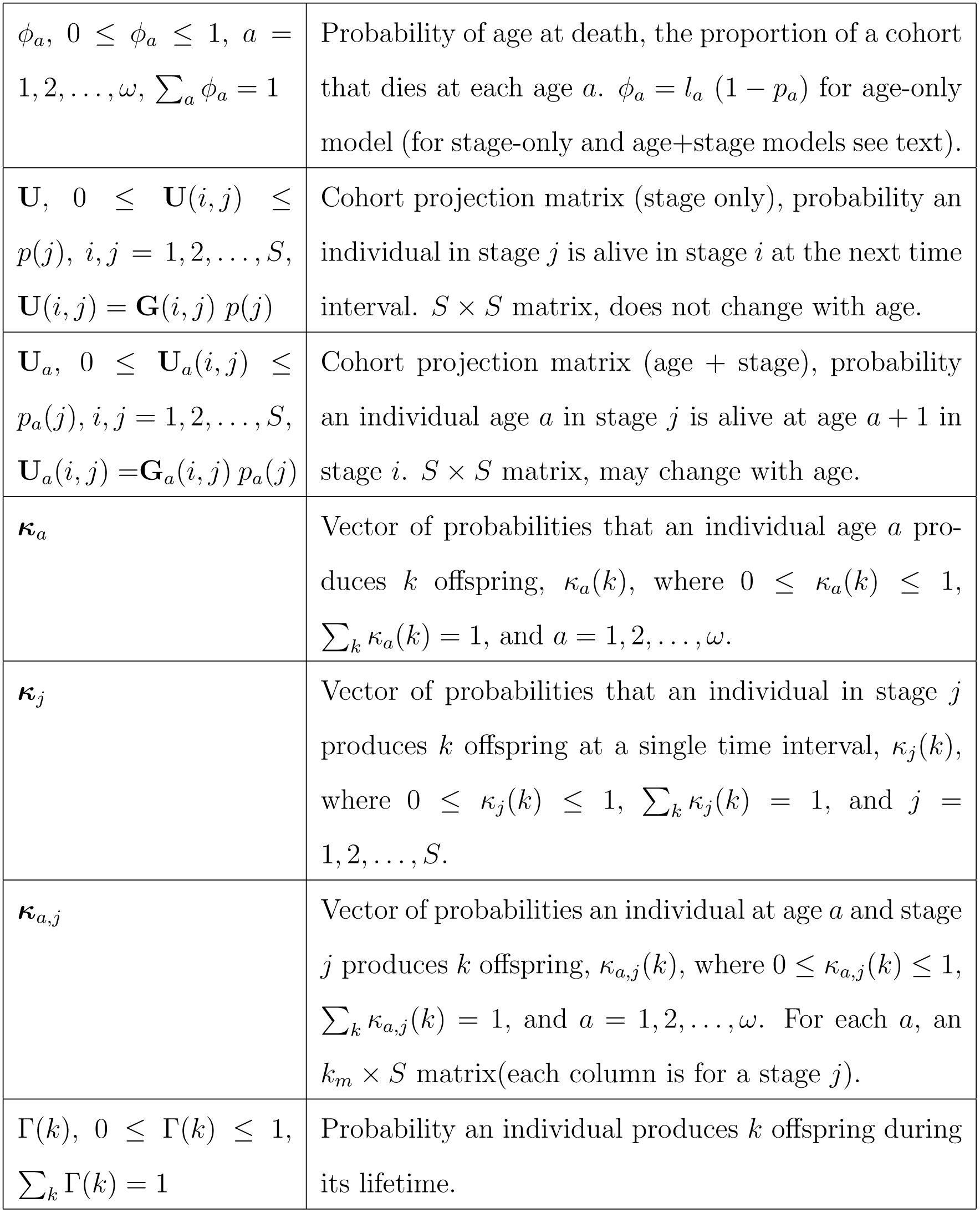
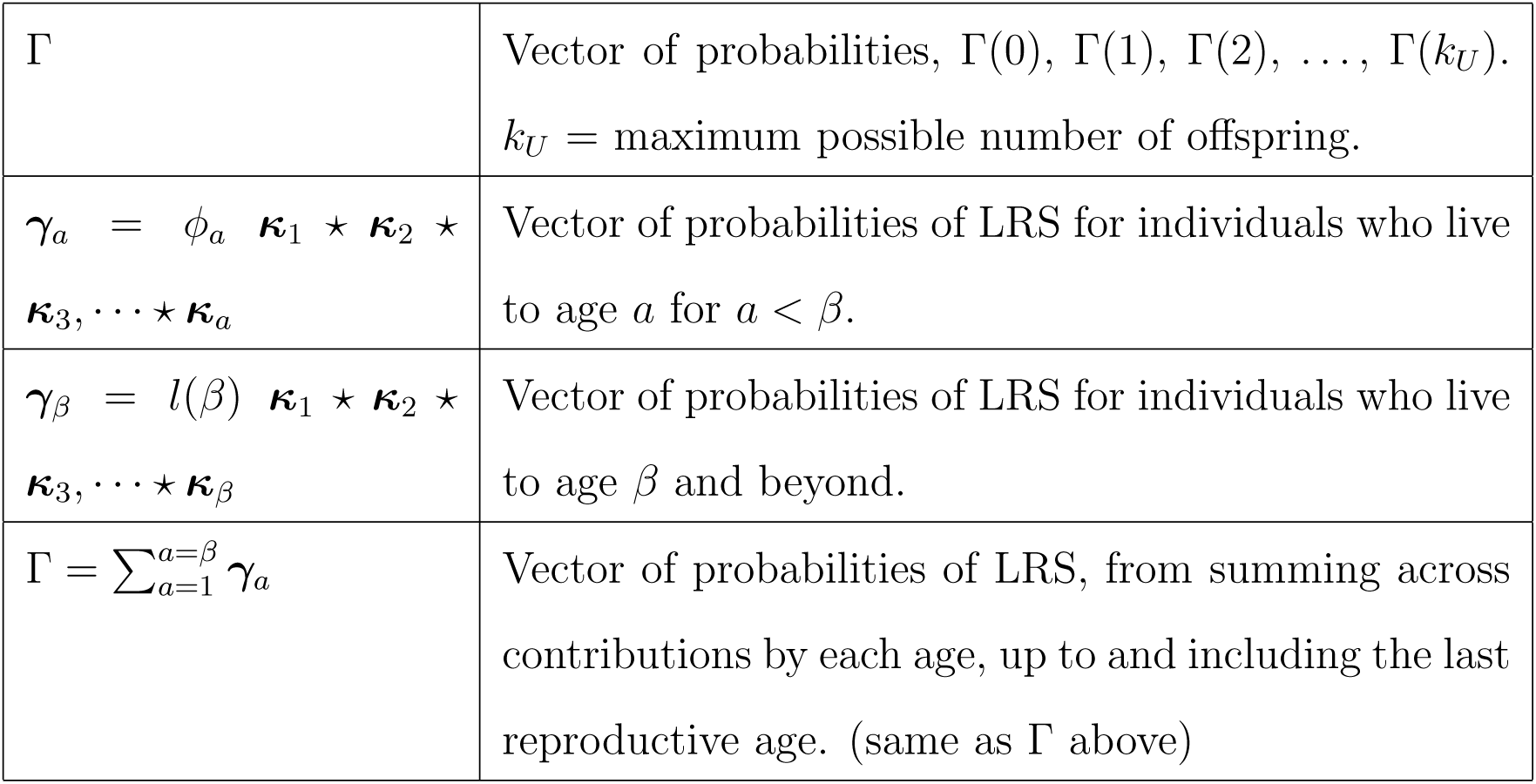
Definitions.

Age-at-death is a random variable (well-defined even for populations where one-period survival depends only upon stage). The probability that an individual dies at each age *a* is the product of the probability of living to the beginning of age interval *a* with the conditional probability of dying during that age interval.

Reproduction for age-only and age+stage models is only possible between some specified youngest age *α* and an oldest age *β*, respectively. Stage-only models may have non-reproductive stages, but such stages may occur at any age. The number of offspring produced during a single time interval is a random variable that can take on integer values *k* = 0, 1, 2, …, *k*_*m*_. The probability distribution of reproduction (in a single interval) depends on age (***κ***_*a*_), stage (***κ***_*j*_), or age+stage (***κ***_*a,j*_), respectively for the age-only, stage-only and age+stage models (Table 1).

Finally, the number of offspring produced by individuals during a lifetime, the LRS, is a random variable that can take on integer values *k* = 0, 1, 2, …, *k*_*u*_. The final goal of our analyses is always the probability distribution Γ whose elements are

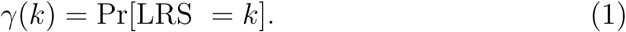

Our calculations assume that, within a single time interval, reproduction precedes death. Thus an individual alive at the beginning of a given time interval always reproduces according to its age and or stage at the beginning of the interval. Our formulas are easily changed to suit other assumptions.

The practical flow of decisions about which of the following methods to use is shown in the online Appendix (Fig. A.4).

### Age-only Model

Here, survival and reproduction during a single time interval depend only upon the age of individuals. We want the probability distribution Γ of LRS (see equation (1)). The distribution of offspring ***κ***_*a*_ at age *a* is taken to be binomial for age-structured human populations (so *k*_*m*_ = 1 and there are 0 or 1 offspring at each age); but our method below applies to any form of probability distribution. We break life into segments: newborns (age 1) who die before reaching age 2, those who live to age 2 but die before 3, and so on. First, consider individuals who die before reaching age *a* = 2. The probability of such a death is *ϕ*_1_ = *l*_1_ [1 *-p*_1_], and reproductive success at age 1 follows the distribution ***κ***_1_, so the LRS for such individuals has the probability distribution

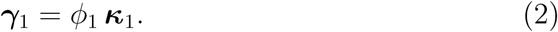

Next, consider individuals who reach age *a* = 2, reproduce and die before age *a* = 3. Such individuals have had two opportunities to reproduce. If these individuals have LRS = 2, they could have: i) produced 0 offspring at age 1 and 2 offspring at age 2, or ii) produced 1 offspring at age 1 and 1 offspring at age 2, or iii) produced 2 offspring at age 1 and 0 offspring at age 2. In other words, at ages (1, 2), they could have (2, 0), (1, 1) or (0, 2) offspring. Hence the probability of LRS = 2 is

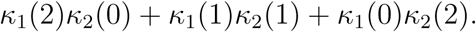

Each LRS = *k* is analyzed in this way, and the corresponding probabilities are elements of the discrete convolution

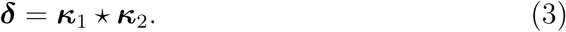

For more detail on convolutions, see the online Appendix (Table A.1 and section A.1). There are (2*k*_*m*_ + 1) = 3 possible outcomes resulting from this convolution. Since the proportion of a cohort that dies at age 2 is *ϕ*_2_, the LRS for individuals in a cohort who die before reaching age *a* = 3 is distributed according to the probabilities

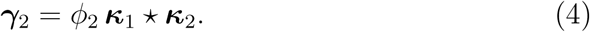

In general then, an individual who lives to age 2 ≤ *a* < *β* but dies before reaching *a* + 1, has LRS distributed as

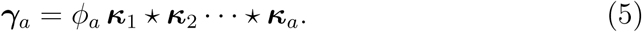

Reproduction ends after age *β*, so survival to later ages does not change the LRS. Consequently, for an individual who lives to age *a* ≥ *β*, the LRS has distribution

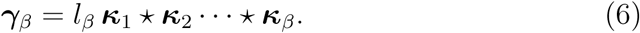

Adding, we have

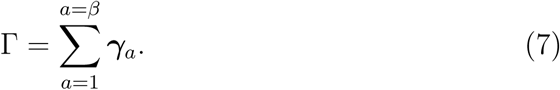

### Stage-only Model

For many species, stage (such as size or developmental stage) is predictive of vital rates rather than age (Lefkovitch 1965). Survival, stage-transition, and reproduction during a single time interval depend only on the stage *s* of individuals at the beginning of that interval.

The number of offspring (*k*) produced during a time interval depends only on stage *j* at the beginning of the time interval and is specified by probability distributions ***κ***_*j*_ = {*κ*_*j*_(*k*)} of producing *k* = 0, 1, 2, …offspring (Table 1). These distributions are often taken to be Poisson (e.g., some cases studied in vDC), but that is not a requirement of our analysis.

Stage-structure requires multiple convolutions that make the age-only procedure computationally unwieldy. This is because here, in contrast to an age-structured model, individuals may visit a stage any number of times before dying. They may also move back and forth between non-reproductive and reproductive stages. So we need a fast way of (a) doing repeated convolutions, and (b) summing over repeated convolutions.

To solve (a) we use the mathematical fact that the (discrete) Fourier transform of a (discrete) convolution is the product of Fourier transforms, and the numerical fact that the FFTW package (Mersmann 2019) does a fast discrete Fourier transform. To solve (b), we exploit our earlier work (Steiner and Tuljapurkar 2012) and combine a closed-form generating function with Fourier transforms, then use an inverse FFT. For the details (or if this brief description makes you curious) see the online Appendix (section A.5).

Thus we can compute the exact distribution of LRS for any stage-structured model. We can separately, and quickly, compute the probability that the LRS = 0. Both analyses are given in the online Appendix (sections A.5 and A.8).

### Age+Stage Model

Here survival, stage transitions and reproduction during a single time interval depend on both age and stage (e.g. Coulson (2012), Table 1 and the online Appendix). In principle, here we can proceed as with the age-structured model. But the computation depends on the “size” of the life cycle, and determines a choice between two options. If all individuals reach a terminal age and die, and if the number of stages is modest, use the age-structured approach. If there are many ages and/or stages, and/or if all individuals reach an “old” age with stages in which they reproduce and remain until death, the age-structured approach is too slow. In that case, use a general stage-structured approach with block matrices (see online Appendix section A.6 for details).

## Results

### Age-Only Model: Two examples

We contrast the Hadza, a hunter-forager group thought to be representative of early humans, to the Hutterites who in the 1950s were famous for their high fertility (vDC). We consider only females with no migration. One-period survival probabilities by age, *p*_*a*_, yield survivorship *l*_*a*_ (the probability of surviving until age *a*). Reproduction ceases at menopause *β* ≈ 50 yr, so we do not need later ages. The commonly used fertility rate *f*_*a*_ is the mean number of offspring at each age. Fertility at age *a* can be 0 or 1 for human populations (vDC), with

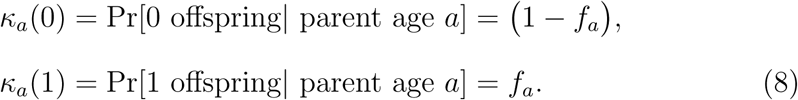

For each age, these numbers constitute a distribution we denote ***κ***_*a*_. We ignore twinning, which is rare in these populations (but can be included by adding the value of *κ*_*a*_(2)). For other species (e.g., if there are litters with many offspring) we need additional information to create a distribution of offspring at each age.

We used our methods to compute the exact distributions of lifetime reproductive success (LRS), Hadza in Fig. 1(a), Hutterites in Fig. 1(b); the distribution is given by the solid points in the figure. Notable features:

**Figure 1:**
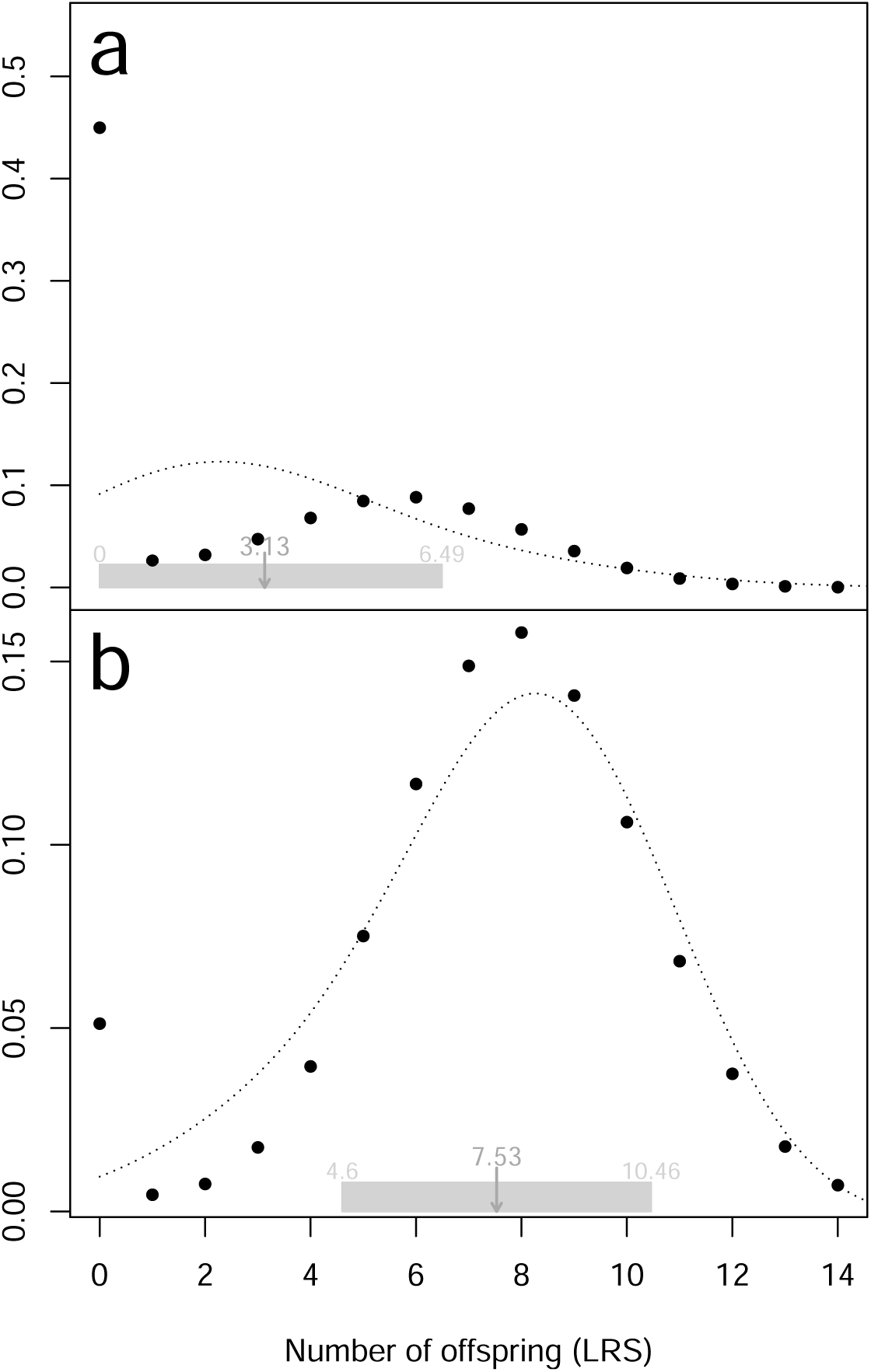
The LRS distribution of Hadza (a) and Hutterites (b) [solid points], and a Gram-Charlier approximation based on the first three moments [dotted lines].

1. The distribution for the Hadza is dramatically bimodal, with a very high mode at 0 and another at LRS ≈ 6. These modes are very different from the mean LRS of 3.13; the spread in LRS is not described by the standard deviation (= 3.36). The third central moment (= 21.7) does imply the distribution is skewed.
2. The Hutterites have a low mode at 0 but a high mode at LRS close to the mean of 7.53. Excluding the mode at 0, the distribution appears to concentrate nicely around the mean value, with a standard deviation (= 2.93) that captures the spread of nonzero values. However the third central moment is negative (= −14.49), presumably because of the mode at 0. If we didn’t have the full distribution on view, this value of the third moment would be hard to understand.

#### Mode at LRS = 0

Is this likely to be a common feature? A female may have 0 offspring if she dies before reproduction begins, or lives on but has 0 offspring at every age. For the Hadza and the Hutterite, reproduction can begin at age 13 yrs. The probabilities of dying by that age are 0.415 for Hadza, 0.043 for Hutterites. The probability of living on but childless is 0.035 for the Hadza and only 0.008 for the high-fertility Hutterites. So the mode at LRS = 0 is largely determined by high infant mortality in the Hadza and the much lower infant mortality among Hutterites. Thus in general we should see a mode at LRS = 0 unless early mortality is extremely low.

#### Do 3 moments predict the distribution?

We can compute the LRS distribution from our methods, but can we recover a reasonable approximation from just the first 3 moments? For the Hadza, the answer is emphatically no, as shown by the Gram-Charlier approximation in Fig. 1(a) (the dotted line). For the Hutterites, the approximation works rather well for nonzero LRS but entirely misses the mode at 0, as shown in Fig. 1(b) (the dotted line); so the answer is still no. We’ve tried a different method, starting with a Poisson and then using a numerical search for the “best” modification, but the answer is still negative. We have not tried using higher moments, because they do not seem likely to change our results.

### Stage-Only Model

#### An example: An evergreen tree *Tsuga canadensis*

We use a stage-only model for *Tsuga canadensis*, eastern hemlock, an evergreen tree that grows to be 40-75 feet tall in the wild (as cited in vDC). Populations of this tree, as noted by Lamar and McGraw (2005), usually include many saplings, small individuals that may remain in good condition beneath a canopy for hundreds of years. So individuals exhibit considerable stasis.

The LRS for *T. canadensis* has an extremely pronounced mode at 0, and the skew of the distribution is so dramatic that we had to plot the logarithms of the probabilities of LRS, as shown in Fig. 2. The average LRS is just 1.42, but the standard deviation of 37.57 is not useful as a measure of spread at either lower or higher values. The third moment is huge, ≈ 2.6 × 10^6^, which certainly means great skew, but does not predict the actual distribution in the figure. We expect a similar skewed distribution for many species (including plants and marine organisms) that produce large numbers of offspring such as seeds or larvae, but few recruits, and even fewer adults. The skew seen in Fig. 2 is relevant to current paradigms about life history evolution and selection, as discussed in the final section.

**Figure 2:**
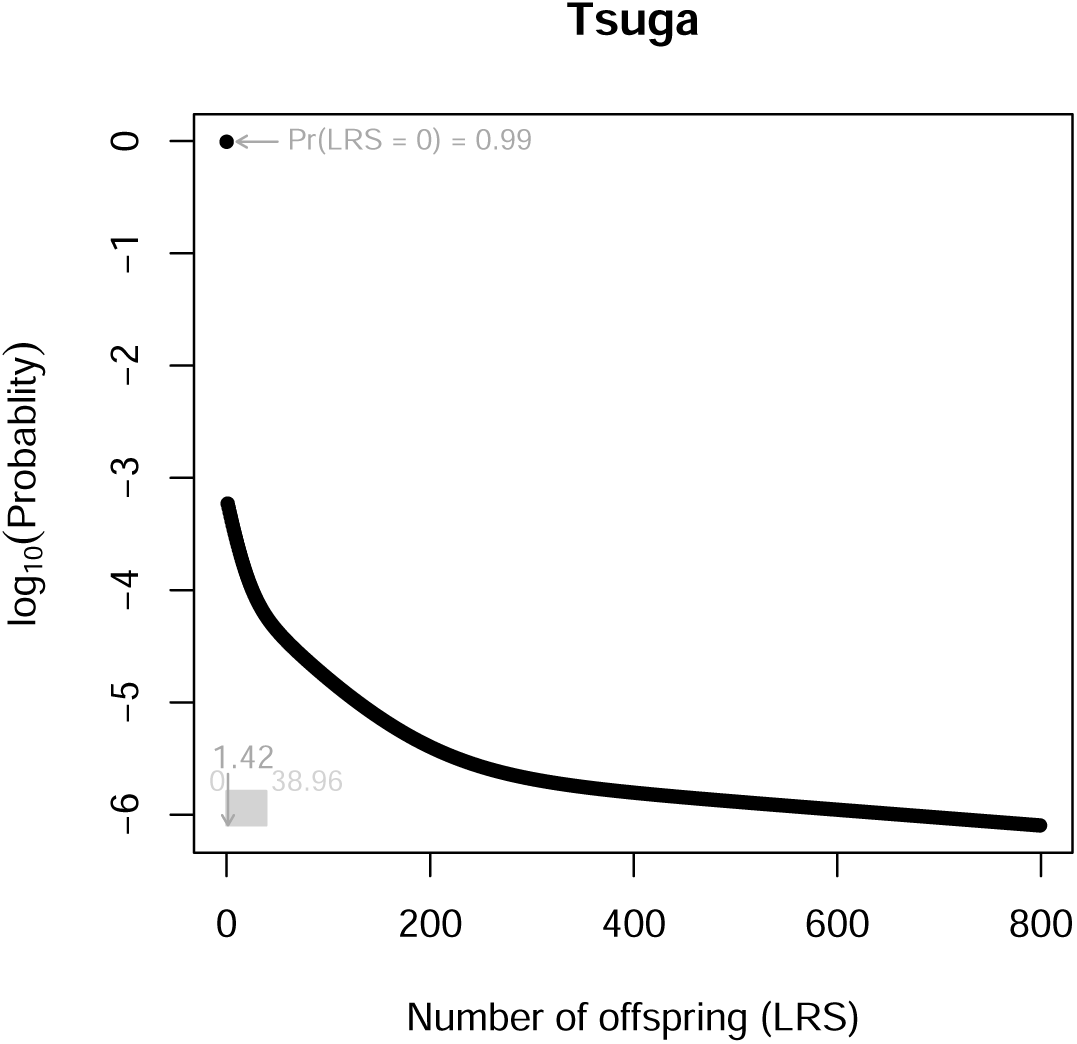
The LRS distribution of *Tsuga canadensis*, eastern hemlock.

#### Mode at LRS = 0

Individuals may produce 0 offspring even when they are potentially reproductive, so our analysis identifies two related but distinct probabilities:

i. One is Pr[LRS = 0], which is a high 0.9883 for *T. canadensis*! This number is the sum of the probability of surviving while in non-reproductive stages plus the probability of producing 0 offspring while in reproductive stages. We can always compute this probability exactly (see online Appendix section A.8).
ii. So does it always make sense to ask (as in the age-structured case) for the conditional probability

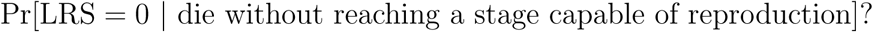 The answer depends on the life cycle. Suppose we start in a non-reproductive stage. In one type of life cycle, every individual that survives eventually makes an irreversible transition to one or more stages that are capable of reproducing (we call these reproductive stages). For this case, it makes sense to treat the reproductive stages as “absorbing,” so we can compute the corresponding absorption probability. Then [1 *-* absorption probability] is just the conditional probability defined above (see online Appendix section A.8). The value of this conditional probability is 0.9878 for *T. canadensis*; comparing this number with 0.9883, we see that most individuals who have an LRS of zero must die without ever having entered a stage that is capable of reproduction. But, in life cycles of stage-structured populations that are not as described above, this probability doesn’t make sense. In the general stage-structured life cycle, individuals can make any number of transitions back and forth between non-reproductive and reproductive stages and death can occur at any point.

### Age+Stage Model

#### An example: Roe deer *Capreolus capreolus*

In general, an individual is classified by both age *a* = 1, 2, …, *ω* and stage *s* = 1, 2, …, *S*. We illustrate with an estimated age+stage model for Roe deer *Capreolus capreolus* (Plard et al. 2015). Age is in years, and size in 200 equal body mass intervals from 1 Kg to 40 Kg. There are 4 age groups; in each group, there is one set of transition and survival rates and reproduction probabilities (the age groups are yearlings, adults age 2-7 yr, adults age 8-11 yr, adults over age 11). Age advances in single years.

The “initial stage” is a combination of age (here, yearlings in age class 1) and size. For initial age *a* = 1 there are potentially 200 initial sizes. In practice, yearling sizes range from size class 40 to 100, but we analyze the full potential size range (see the online Appendix section A.10). Fig. 3 plots the LRS distribution for 3 different initial size classes, 1, 75, 200. The distribution shifts from unimodal at LRS = 0 for the smallest yearlings, to unimodal at a nonzero LRS for the largest yearlings, and can be multimodal at intermediate sizes.

**Figure 3:**
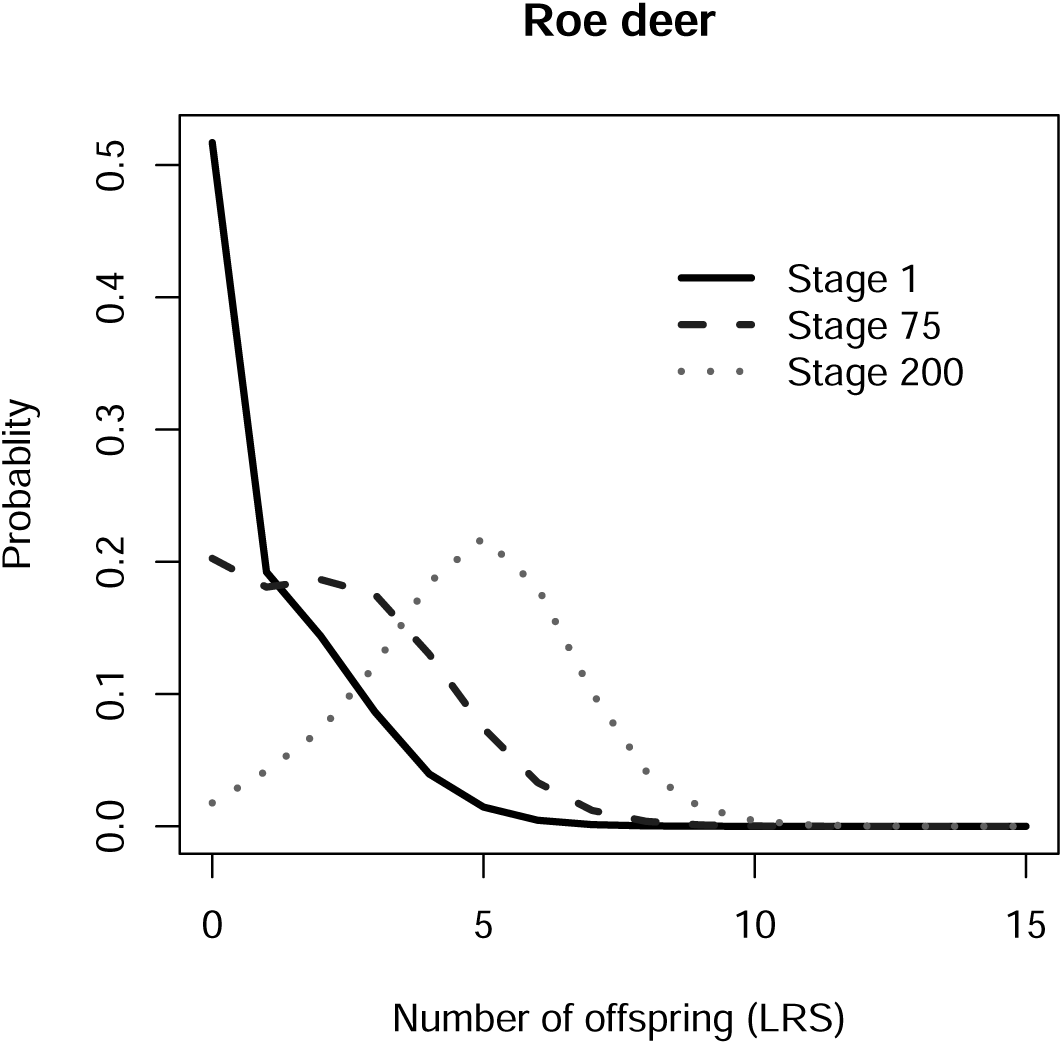
The LRS distribution of yearlings of Roe deer *Capreolus capreolus* for different initial size classes of 1, 75 or 200.

Very small offspring have a high mode at 0, and thus the highest probability of leaving no offspring – no surprise. Neither the average LRS nor the standard deviation seem to be useful descriptors of the LRS distribution for the smallest yearlings (plotted in the online Appendix Section A.10). Somewhat bigger yearlings (birth size class 70-84, see Fig. 3) have a bimodal distribution of LRS. Larger juveniles (size class 85 upward) have a single nonzero mode but still high odds of leaving 0 offspring. The nonzero mode shifts from an LRS ≈ 3 (for size class 80) to an LRS ≈ 5 (for size class 200) with increasing yearling size, and the distribution of LRS concentrates towards the mode. Very large offspring have a high nonzero mode at LRS ≈ 5, with a normal-like LRS distribution around the mode.

#### Mode at LRS = 0

The height of a mode at an LRS of 0 can be computed using a age+stage method described above (see online Appendix Section A.10). We found that Pr[LRS = 0] declines with increasing birth mass. This is expected, since in this population it pays to be born larger. However the gain from increasing birth weight is strikingly nonlinear, so that the advantage of being born 1 Kg heavier is much greater for small than for large yearlings.

To dissect this effect, we analyzed yearling death probability and the probability of having no offspring if you live to later ages (see online Appendix). There is a drop in both as birth mass increases, but the gain in yearling survival reduces faster than the gain in later survival/reproduction.

## Discussion

We have presented an analytical method that computes the probabilities of getting every possible LRS, 0, 1, 2, 3, …. We illustrated our results for two human populations for age-structure, eastern hemlock for stage-structure, and Roe deer for age+stage structure. We found that the LRS distribution often has one mode at LRS = 0, and a second at nonzero LRS. For several of our examples, the mean, variance and third moment are poor descriptors of the LRS distribution. We hasten to add that these moments are useful descriptors if offspring (often censused as newborns or recruits) have high survival, probably uncommon in nature but common in modern human populations. Even in the latter case, though, we recommend using our methods to analyze the probability that LRS = 0. We find an even greater variability of the LRS than expected from previous work (Tuljapurkar et al. 2009b; Steiner and Tuljapurkar 2012; van Daalen and Caswell 2015; Snyder and Ellner 2018) and vDC. Here we discuss extensions and implications of our results, and directions for future work.

### Descriptors and Sensitivity analyses

How best to describe the variability that we have found in the LRS? And given a descriptor, what are its sensitivities to vital rates? In ecology, sensitivities determine how vital rates (or underlying parameters) affect the LRS distribution, whereas in evolution sensitivities capture selection pressures that depend on LRS Caswell et al. (2018). For future analytical work, note that our stage-structured analysis applies to any age+stage combinations.

One descriptor of the inequality of LRS is a Gini index (Weiner and Solbrig 1984; Shkolnikov et al. 2003), which corresponds to the mean of absolute differences in individual LRS relative to the average LRS, and is bounded between 0 and 1. The Gini index is commonly used in economics and social science to quantify inequality, but less frequently in ecology and evolution. It has been used to assess variation in the distribution of reproductive success of parasites (Dobson 1986), in the mass and fecundity distributions of a cestode infesting pikes (Shostak and Dick 1987), in the strength of forest edge effects (Matlack 1994) and in size hierarchy studies of competitive processes in plants (Weiner and Solbrig 1984). It has also been recently used to describe the shape of mortality within a population (Archer et al. 2018) and to quantify species differences in the distribution of reproductive ages across vertebrates (Healy et al. 2019). We propose that it will be valuable in analyzing variation between sexes or across species in the LRS distribution. To compute the Gini index, find the cumulative distribution

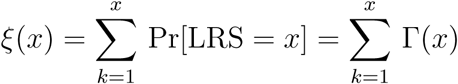

and plot *ξ*(*x*) versus values (*x/k*_*U*_) of the LRS (remember *k*_*U*_ is the largest possible LRS). This is the Lorenz plot shown for our examples in the panels of Fig.4. Equality means that every individual has the same odds of any LRS and is indicated by the dotted lines. The actual distributions are shown by the solid lines and indicates that a few individuals do most of the reproducing. The Gini index (shown on the plots as *GI*) is

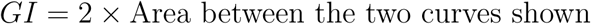

**Figure 4:**
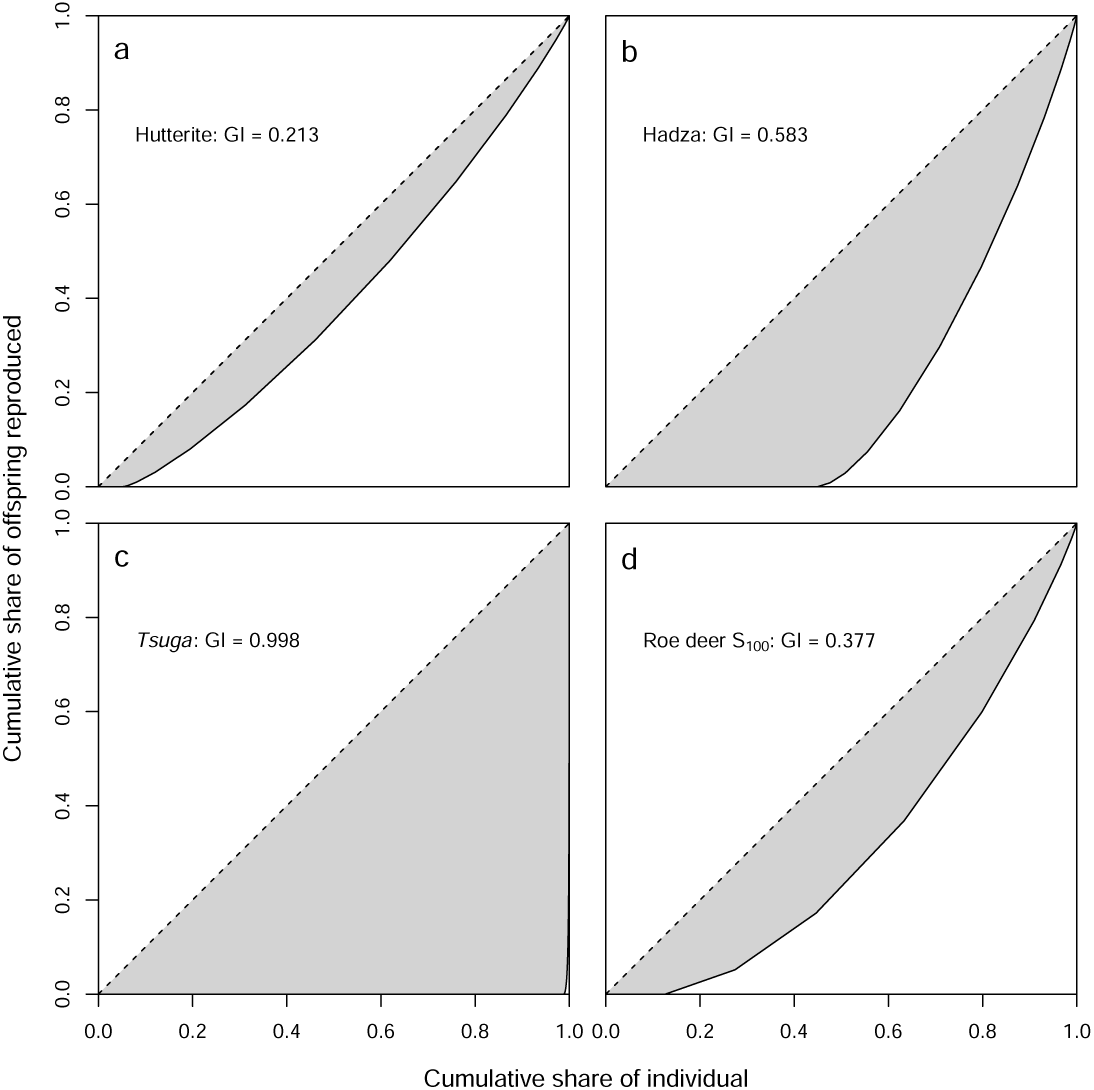
The Lorenz plots for Hutterite, Hadza, *T. canadensis*, and Roe deer (with initial stage 100).

Equality means that *GI* = 0 while complete inequality means that *GI* = 1. As we expect from Fig. 1, the Gini is much smaller for Hutteries than the Hadza. And the extreme inequality of LRS for *T. canadensis* is reflected in a very high Gini. Surprisingly (to us) the LRS for a fairly large Roe deer yearling (size class 100) is also quite unequal, with a Gini value halfway between Hazda and Hutterite.

A general approach to the sensitivity of inequality in LRS should be feasible using our results in conjunction with the methods in van Raalte and Caswell (2013) and Caswell et al. (2018). More generally, numerical perturbations can be used to examine changes in Γ produced by changes in vital rates, and we can compare distributions using, e.g., the Kullback-Leibler distance (Cover and Thomas 2012).

A useful measure highlighted by our examples is the probability of having no reproductive success:

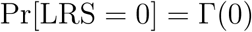

A general expression and computational method for this probability is given in the online Appendix (section A.8). Derivatives of this probability with respect to these parameters should be computable using known approaches, e.g., Caswell et al. (2018) or Steiner et al. (2012).

Another useful measure is the nonzero modal value, call it *k*^*^, of the LRS – if it exists. We found a nonzero mode in several examples, but there is no nonzero mode for the tree (*T. canadensis*). Assuming its existence, a nonzero mode means that

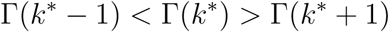

This modal value may be more meaningful in some populations than the commonly used average LRS. We can use our result to find *k*^*^ and the methods in Caswell et al. (2018) should be extendable to compute sensitivities.

### Implications

In our age-structured applications each individual is governed by the same set of demographic rates (probabilities). In other words, all individuals have the same chances of surviving and reproducing at each age. Yet, the distribution of LRS is multimodal; this reveals how much individual heterogeneity can be generated by chance alone. This variation caused Steiner and Tuljapurkar (2012) to conclude that entirely neutral processes could generate the empirically-observed multimodal distributions of LRS so frequently reported.

In our stage and age+stage structured models individuals that start life with different phenotypic trait values can have contrasting distributions of LRS. For each phenotype there is still a large amount of individual heterogeneity in LRS similar to that observed for our age-only models, but for individuals with different phenotypes we will find differing LRS distributions. These differences will generate opportunities for selection. Whether such selection would result in evolution will depend upon whether there is a genetic basis for individuals entering the population with different phenotypes.

Stage- and age+stage structured models are powerful. They can be constructed for stochastic, density-dependent, and frequency-dependent environments to predict population (Tuljapurkar 1990), life history (Tuljapurkar et al. 2009a; Caswell 2001), and evolutionary dynamics (Coulson et al. 2011, 2017; Childs et al. 2016). A large number of ecological and evolutionarily important statistics can be calculated from these models, with analysis of these models revealing how different values of these statistics arise. By extending the toolkit of structured models to predict entire distributions of LRS we open the door for a number of powerful, new analyses. For example, we can ask how different growth trajectories will impact distributions of LRS, we can gain insight into why weak associations between birth weight and LRS are so frequently observed, and the conditions when lifetime measures of reproduction act as a good, or bad, surrogates of fitness. In addition, we could use our approach to generate distributions of other per-generation life-history statistics such as generation times, or future life expectancies at a given age.

Previous analyses of LRS distributions in vertebrates relied on the standardized variance and have supported the existence of a link between mating systems and sex differences in LRS distributions, with similar distributions in males and females of monogamous species and much more widespread and skewed distributions in males than in females in highly polygynous species (Andersson 1994). However, our findings demonstrate that only using the first few moments such as mean and variance does not capture important characteristics of an LRS distribution. More comprehensive metric such as the Gini index are much more appropriate and should be used in future comparisons of LRS distributions. Our approach should thus allow accurate comparisons among LRS distributions that are currently accumulating thanks to long-term individual-based monitoring of populations in a broad range of species across the tree of life. Moreover, in addition to sex and species, our approach provides a suitable way to assess how ecological factors (e.g. environmental harshness), life history tactics (e.g. slow vs. fast life cycle, capital vs. income breeding), and population structure (e.g. in terms of age, size) influence LRS distributions within and among populations.

The ability of the discrete-time structured model to gain ecological, life history, and evolutionary understanding is remarkable, and with each year new uses for them are developed. Such utility is not surprising, as these models (Coulson et al. 2011) include the fundamental biological processes of development, inheritance (although not considered in the current paper), survival and reproduction, and these processes can be specified to vary across individuals as a function of age, sex, phenotype, genotype (although not considered in the current paper) and environment. It is consequently unsurprising that analysis of such models has generated so much understanding in recent years. The methods we develop here open up opportunities for yet more biological insight.

## Acknowledgements

We thank Ni Lao, Hal Caswell, and Maarten Broekmann for useful comments.

## Appendix

### A.1 Discrete Convolutions and Age structure

Consider an age-structured population with only two age classes. Say that an individual of age class 1 has some number of offspring *i*_1_ = 0, 1, …, *k*_*m*_ with probabilities ***κ***_1_ = {*κ*_1_(*i*_1_)}, and an individual of age class 2 has *i*_2_ = 0, 1, …, *k*_*m*_ offspring with probabilities ***κ***_2_ = {*κ*_2_(*i*_2_)}. Now consider an individual who may produce offspring at both ages 1 and 2, and call *δ*(*n*) the probability that such an individual produces *n* offspring, *n* = 0, 1, 2, …, 2*k*_*m*_. Given any *n*, some *m*_1_ offspring are produced at age 1 and the rest, *k*_2_ = (*n* − *m*_1_), must be produced at age 2. The number of ways to obtain a given *n* varies with *n*. For example, there is only one way to obtain *n* = 0 (both *m*_1_ = 0 and *m*_2_ = 0). Similarly, there is only one way for *n* = 2*k*_*m*_ (both *m*_1_ = *k*_*m*_ and *m*_2_ = *k*_*m*_). There are only two ways for *n* = 1 (either *m*_1_ = 1 and *m*_2_ = 0 or *m*_1_ = 0 and *m*_2_ = 1). In general, we sum over the probabilities of the different ways we could get a particular *n*

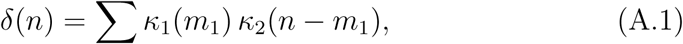

where the number of terms in the summation varies from 1 to a maximum of *k*_*m*_ + 1. The probabilities {*δ*(*n*)} are components of a distribution that we write as ***δ***. Also, the fact that the first argument in equation (A.1) is *m*_1_ and the second is (*n - m*_1_) means that ***δ*** is what is called a discrete convolution of the distributions ***κ***_1_ and ***κ***_2_, written as

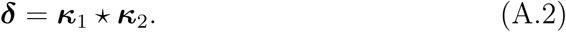

Alternatively we can describe ***κ***_1_ and ***κ***_2_ by the probability generating functions

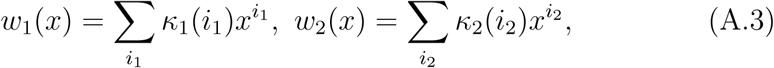

where *x* is a dummy variable. Let the probability generating function for ***δ*** be

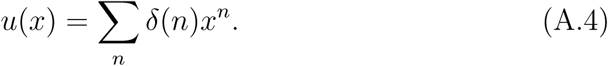

Setting

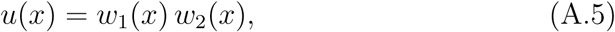

also (algebra!) yields the probabilities in equation (A.1). Thus there is a correspondence between convolutions of distributions as in equation (A.2)) and products of generating functions (as in equation (A.5)).

Using equation (A.3), we see that the distribution ***κ***_1_ ⋆***κ***_1_ has a generating function [*w*_1_(*x*)]^2^. As a result, we write ***κ***_1_ ⋆ ***κ***_1_ as ***κ***^2^. Also, ***κ***_1_ ⋆ ***κ***_2_ = ***κ***_2_ ⋆ ***κ***_1_. Finally, note that convolutions can be rapidly computed (for discrete and continuous functions).

For age structured populations, we can use repeated convolutions to compute the LRS distribution as in the main text. But this method doesn’t work for stage structure so we need a different approach that starts with the property discussed next.

### A.2 Fourier transforms

Fourier transforms require complex numbers. Denote

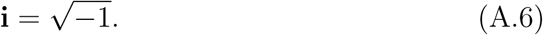

For an integer *N*, define the frequencies

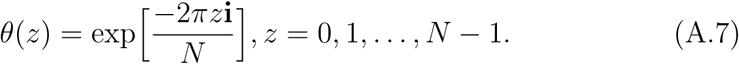

Then for any integer *q*, define the discrete function ***κ*** = {*κ*(0), *κ*(1), *κ*(2), …, *κ*(*q*)} (this could be a probability distribution). Then we define the Discrete Fourier Transform (DFT) of ***κ*** as a function 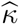, whose values are given for each frequency in equation (A.7) as,

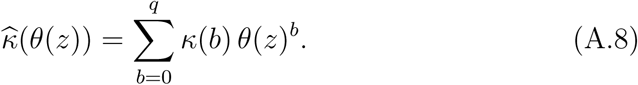

Now suppose that ***ψ*** is another series of numbers, ***ψ*** = {*ψ*(0), *ψ*(1), …, *ψ*(*q*)}. For frequencies *θ*(*z*), we can use equation (A.8) to find the Fourier transform 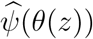. It is well known that (James 2002)

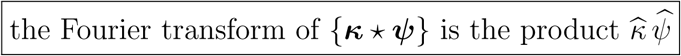

In consequence, for any frequency *θ*(*z*) in equation (A.7): for any integer *m* ≥ 0,

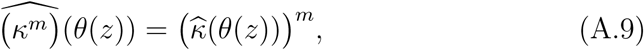

and for any non-negative integers *m*_*a*_, *m*_*b*_

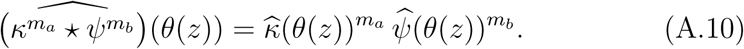

### A.3 Digression: A Generating Function

We consider a general age+stage model following Steiner and Tuljapurkar (2012). Age and stage are used together: given *A* ages and *S* discrete stages, there are (*A* × *S*) = *S*_1_ unique combinations, so that we can proceed as if we had a stage-only model with *S*_1_ stages. Let there be *S*_1_ living stages, with transitions between them with probabilities **U** (Table 1 in main text). Form the matrix **U**^*T*^ = **Q**. The element *q*_*ij*_ of matrix **Q** is the probability of being in stage *i* in one time interval and then surviving to reach stage *j* in the next time interval. The stages that are rows (or columns) of **Q** are transient, meaning that an individual starts in any stage then makes transitions to other stages until an eventual but certain death. Starting in a particular stage *i* let *τ*_*j*_ be the (random) time spent in stage *j* before death. (In the main text we use matrix **U**, but here, we are using the transposed matrix **Q** and so reverse rows and columns).

The death probabilities for stages 1 to *S* are components *d*_*i*_ of a vector **d**. These death rates are of course simply probabilities of not reaching another transient stage, so e.g.,

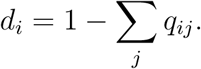

We summarize by defining

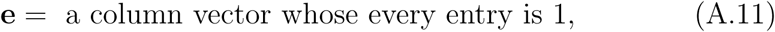

and writing **I** for the identity matrix, so that

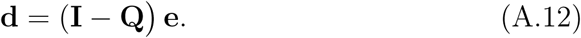

Consider the possible outcome in which an individual spends a time *m*_*j*_ in stage *j*. In our notation, this outcome means that *τ*_*j*_ = *m*_*j*_, conditional on starting in stage *i*, and has probability,

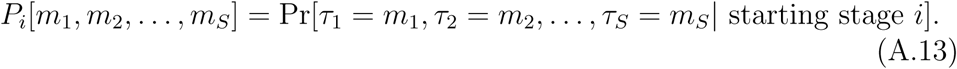

Let *w*_*i*_, *i* = 1, …, *S* be a set of (dummy) variables, and make a generating function for the above probabilities,

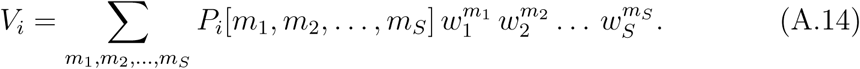

Say we start in stage 1, so *i* = 1. Then *V*_*i*_ = *V*_1_ is a polynomial, whose first term is *w*_1_*d*_1_ – meaning an individual starts in 1 but dies before making a transition. The second term of *V*_1_ is

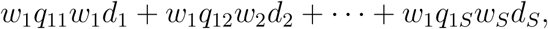

meaning an individual starts in stage 1, lives one period and reaches some stage *j*, but then dies before making a second transition. And so on.

Following Steiner and Tuljapurkar (2012), form the diagonal matrix

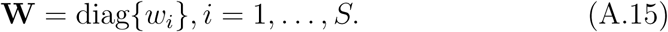

Define the vector function

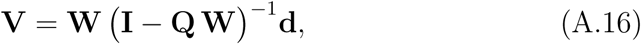

where **I** is the identity matrix and **d** is defined in equation (A.12).

Let the components of **V** in equation (A.16) be {*V*_*i*_, *i* = 1, …, *S*}. Then this component *V*_*i*_ is indeed the generating function defined earlier in equation (A.14). The reader can derive this by expanding the inverse in equation (A.16) and seeing, e.g., that the resulting polynomial *V*_1_ has the same terms as we describe above.

### A.4 To the LRS

We now combine convolutions, the generating function, and Fourier transforms.

Start by defining frequencies. Choose a number *N* that is the largest possible number of offspring. This may be approximated by a product like *k*_*U*_ ≤ (*ω* × *k*_*m*_) with *ω* being an estimated maximum age, and *k*_*m*_ being the maximum number of offspring per time period. Or a biological upper limit can be used. Either way, we can test the adequacy of our choice by computing Γ and then its sum, which should be very close to 1.

Given *N*, for every stage *i* we pad each offspring probability distribution ***κ***_*i*_ by adding zeros so that the padded distribution has *N* elements. For each frequency *θ*(*z*), as in equation (A.7), let the Fourier transforms of the padded distributions ***κ***_*i*_ be 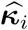, so for example

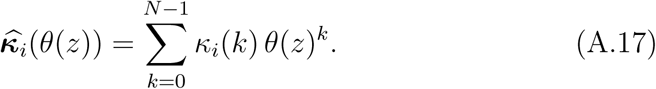

For any non-reproducing stage *i* we have *κ*_*i*_ = {1, 0, …}, and so

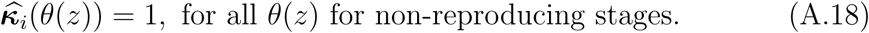

Now as in the previous subsection, say that, starting in stage *i* an individual spends time *m*_*j*_ in each stage *j* before death (for stages *j* = 1, 2, …). The total offspring numbers that result have the distribution

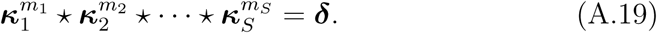

So in this case

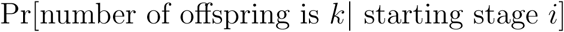

is given by the *k*th element of the convolution *δ* in equation (A.19)

The lifetime probability of having *k* offspring is the *k*th element *γ*(*k*) of

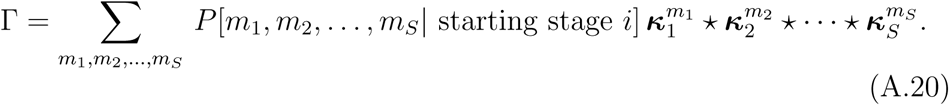

Here Γ can be thought of as a function of the discrete argument *k* = 0, 1, …with values *γ*(0), *γ*(1), …. This is reminiscent of equation (A.14), but unfortunately here we have convolutions and not products of numbers. This is where Fourier transforms come in.

We denote the Fourier transform of this function by 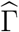. Note that this latter transformed function is defined for each discrete frequency; so if the frequencies are *θ*(0), *θ*(1), …, the Fourier transform is the set of (scalar) values 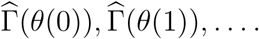

Using the facts about Fourier transforms and equation (A.20) we conclude that

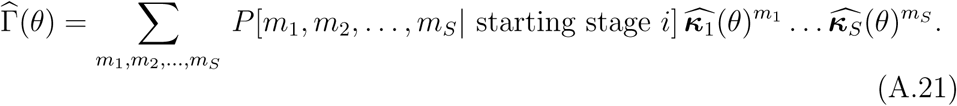

Now this is indeed similar to equation (A.14), so the sum here can be exactly computed using the generating function equation (A.16).

### A.5 Method for Stages

Start with choosing a large number *N*, fix the starting stage *i*, then find the frequencies as in equation (A.7). For each frequency *θ*(*z*) and every stage *j*, compute the Fourier transforms 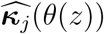, and then make the matrix

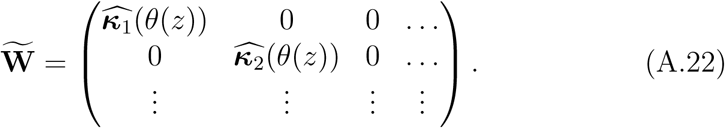

Remembering the generating function, use equations (A.21) and (A.22) to compute the quantity

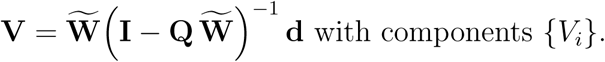

Then use equation (A.21) to conclude that

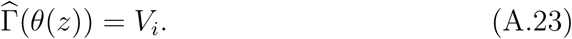

Repeat for all frequencies. Thus one computes the set of (scalar) values 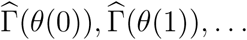 and hence the function 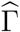.

Final step: use the Inverse FFT to find Γ.

Note: since every age+stage model can be cast as a stage-only model, this method can also be used for age-only or age+stage models. But the age-structured analysis in the main text explains the basic logic that the general method does not. The age-based method is often computationally useless for stage based or age+stage models when we have a large number of stages, or when individuals can stay in a stage for a long time. In such cases, the method here is essential.

### A.6 Block method for age+stage

An age+stage model has ages *a* = 1, 2, …, *ω*, stages *s* = 1, 2, …, *S*. A unique age+stage combination is written *a, s*, and there are *A* × *S* such combinations. In some cases, the general method of the preceding section may be computationally lengthy and the method described below is faster.

#### A.6.1 Inverse

To make it efficient to find the generating function, start by computing the inverse of the matrix

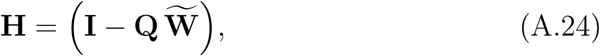

in which **Q** = **U**^*T*^. Here 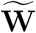 is a diagonal matrix as in equation (A.22). Suppose that we have *S* stages at each age, and that ages *a* = 1, 2, …, *A -* 1 have corresponding and distinct *S* × *S* transition matrices 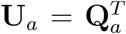. The transition matrix 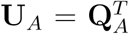 for age *A* is repeating, and applies at age *A* + 1, *A* + 2, …until death. Note that for this case, there are total *A* * *S* stages. Write the diagonal elements of 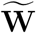 for a given frequency, *θ*(*z*),as

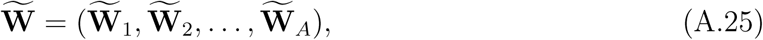

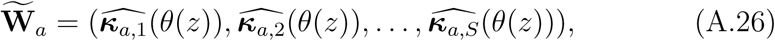

and the matrix of transition probabilities as

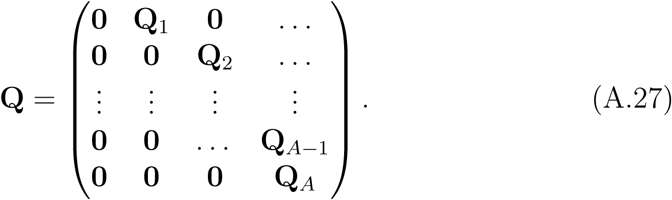

We can write the matrix of equation (A.24) in block form as

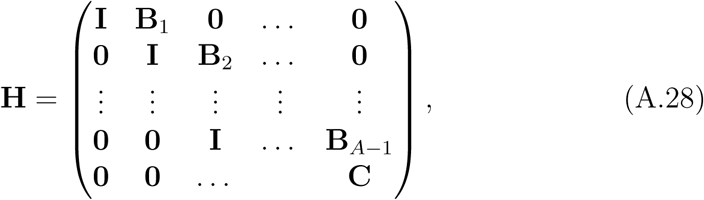

where

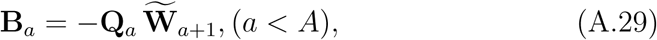

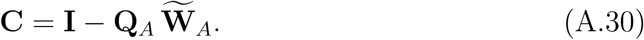

Now use the result in Singh (1979) applied to block matrices. It is best to write this first in cases and then in general. First, say that *A* = 5 so that

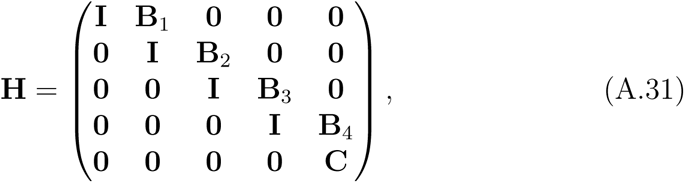

where

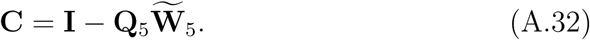

Then

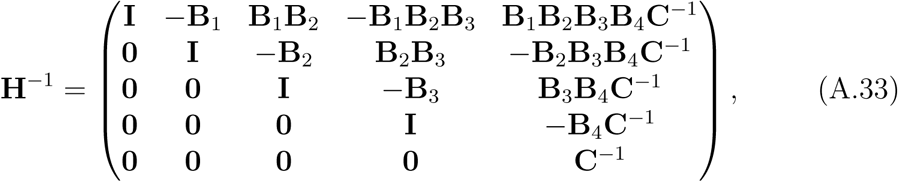

In general, the inverse matrix for equation (A.28) has the form

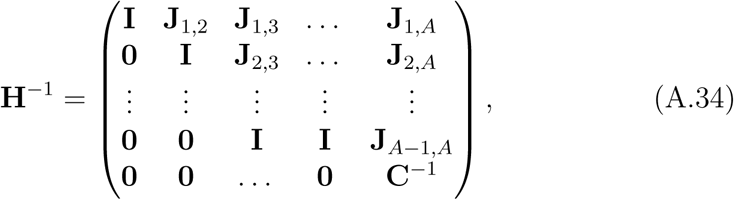

Here,

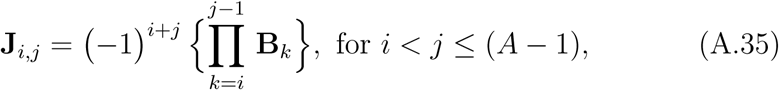

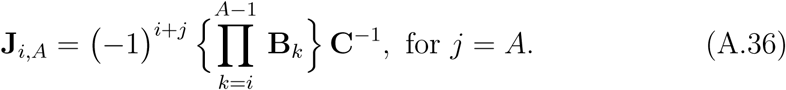

#### A.6.2 The Generating Function

Fix a starting stage *i* out of all *A* × *S* combinations, where *S* is the number of stages in a given age and *A* is the age where the transition matrix **U**_*A*_ starts repeating for following age. Death rates by age+stage are elements (there are *A* × *S* elements) of the vector

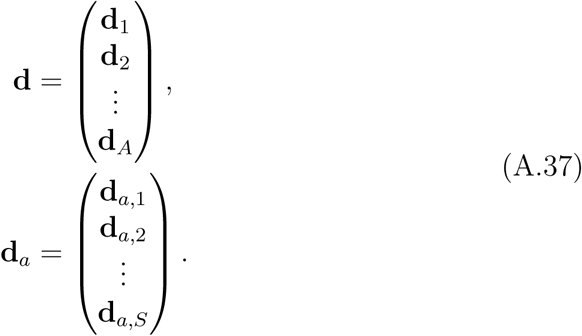

The generating function is

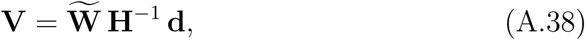

We start with the product

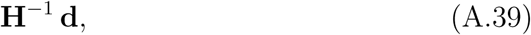

which is a vector of *A* blocks, each *S*-long.

Using the pattern in eqations (A.33 – A.36), we define a sequence:

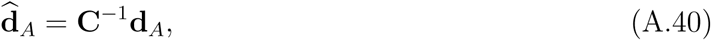

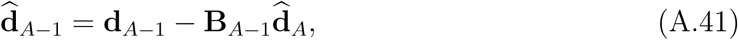

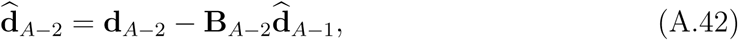

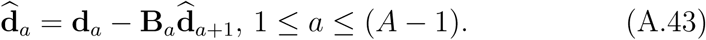

Then set

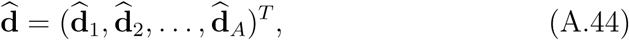

so finally

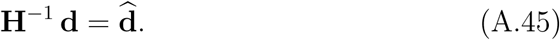

Therefore the generating function is

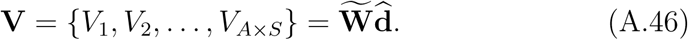

This quantity gives us the Fourier Transform 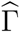 of the LRS distribution Γ – which is the distribution we want. So at the given frequency *θ*(*k*) we have found that

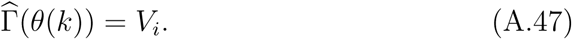

Repeat for all frequencies. Thus one computes the set of (scalar) values 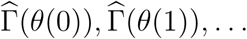 and hence the function 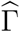. The inverse Fourier transform then yields Γ.

### A.7 Special probabilities: Ages only

To compute Pr[LRS = 0], observe that an individual has 0 offspring if it either (a) dies before reaching reproductive age *α* or (b) survives through the pre-reproductive period, but has 0 offspring thereafter. At reproductive ages *a* the probability of having 0 offspring is *κ*_*a*_(0). Assuming survival, the cumulative probability of having had no offspring by age *a* is

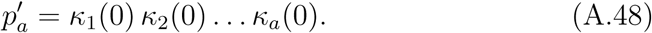

Thus,

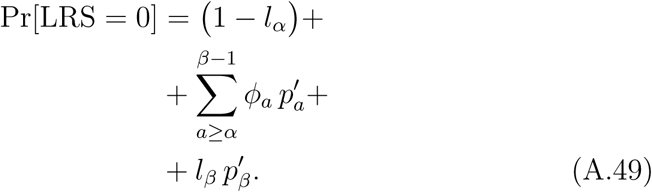

The first line gives us (a) above, the second line adds in reproduction at intermediate ages weighted by the odds of death, and the third line gives us the final contribution to childlessness.

### A.8 Special probabilities: Stages

To compute Pr[LRS = 0] here, we need a different method. Each stage *i* has a corresponding probability that an individual in that stage produces 0 offspring. This probability is 1 for non-reproducing stages, and *κ*_*i*_(0) for all other stages *i*. Define **W**_zero_ to be a diagonal matrix whose elements are

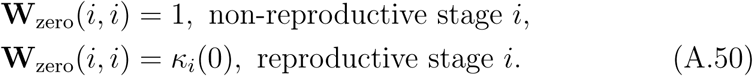

Use the definitions of the preceding section and compute

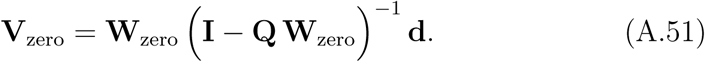

Then the *i*th component of **V**_zero_ is

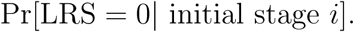

Since every age+stage model can be cast as a stage-only model, this answer can also be used for any model.

For some life cycles we can also compute

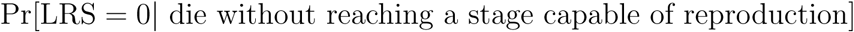

Suppose we start in a non-reproductive stage. In suitable lifecycles, every individual eventually makes an irreversible transition to one or more reproducing stages, or dies. Here it makes sense to treat the reproducing stages as “absorbing” and then (1 - the absorption probability) is just what we want. See Caswell (2001) or Kemeny and Snell (1976) for details on how to do this.

### A.9 Special distributions

#### A.9.1 Poisson

Here the average fertility is *f*_*i*_ for stage *i*, and ***κ***_*i*_ is a Poisson distribution. So ***κ***_*i*_ has the probability generating function

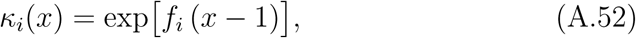

for *x* a dummy variable.

So for stage *j*, and any frequency *θ*(*k*) as defined in equation (A.7), the Fourier transform is

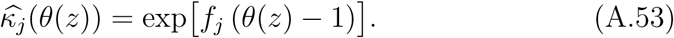

#### A.9.2 Binomial

Here the average fertility is *f*_*i*_ for stage *i*, and ***κ***_*i*_ is a Binomial (with 1 trial, also known as a Bernoulli) distribution. So ***κ***_*i*_ has the probability generating function

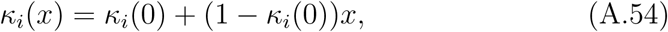

where *κ*_*i*_(0) is the probability that an individual in stage *i* has 0 offspring and *x* is a dummy variable.

So for stage *i*, and any frequency *θ*(*k*) as defined in equation (A.7), the Fourier transform is

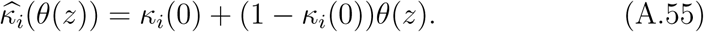

### A.10 Roe deer plots

Small offspring have a high mode at 0, and thus the highest probability of leaving no offspring. Neither the average LRS nor the standard deviation seem to be useful descriptors of the LRS distribution for the smallest yearlings (plotted in the Appendix, Fig. A.1). For the full spectrum of size class, Fig. A.2 shows 41 initial stages spread over size class 1 to 200. Between size class 5 to 200, there are 5 size class interval between each curve. Fig. A.3 shows the Pr[LRS = 0] declines with increasing birth mass.

**Figure A.1:**
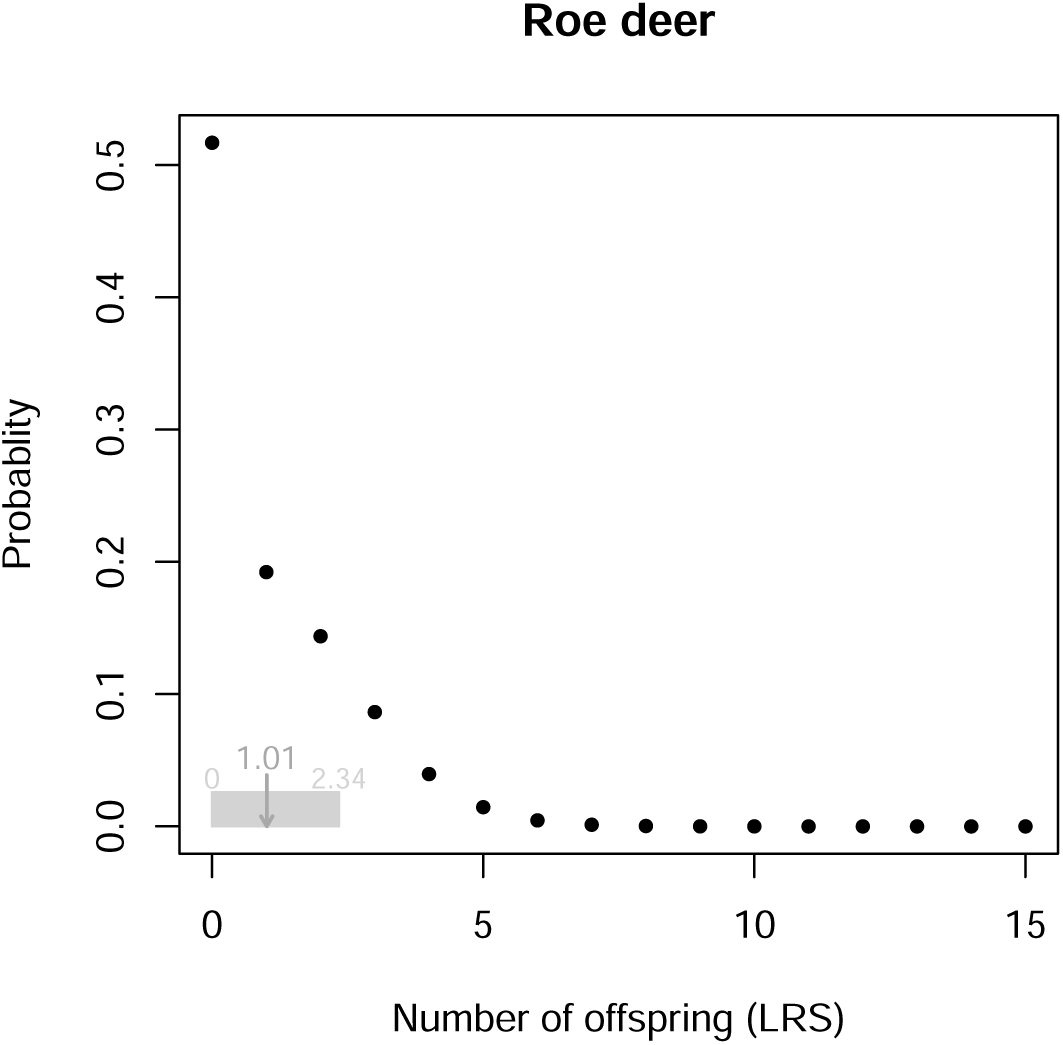
The LRS distribution of Roe deer *Capreolus capreolus*. Mean LRS = 1.01 and the standard deviation is 1.34.

**Figure A.2:**
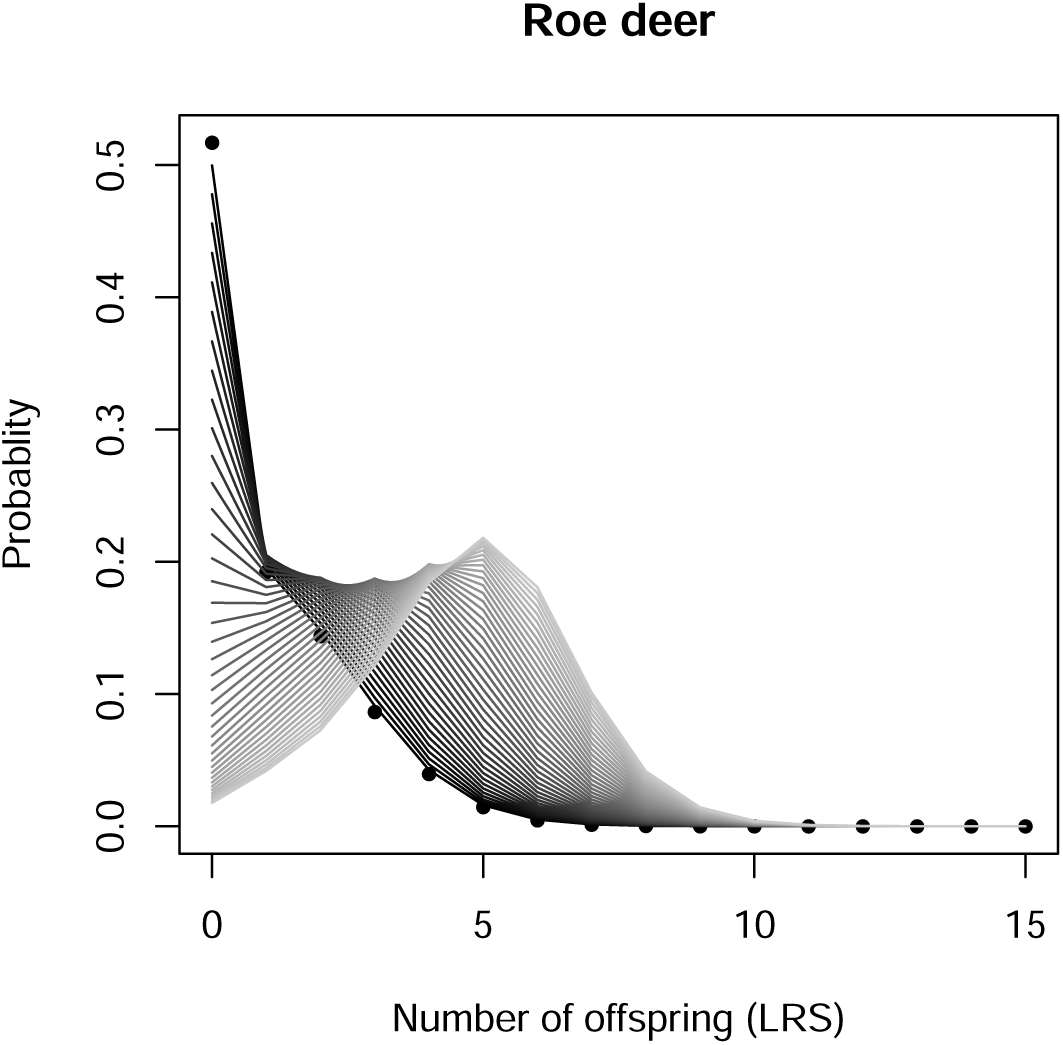
The LRS distribution of Roe deer *Capreolus capreolus*. There are 41 initial stages. Solid points are for yearlings in size class 1. The gradient gray lines are the distribution spread over size class 5 to 200 with interval 5.

**Figure A.3:**
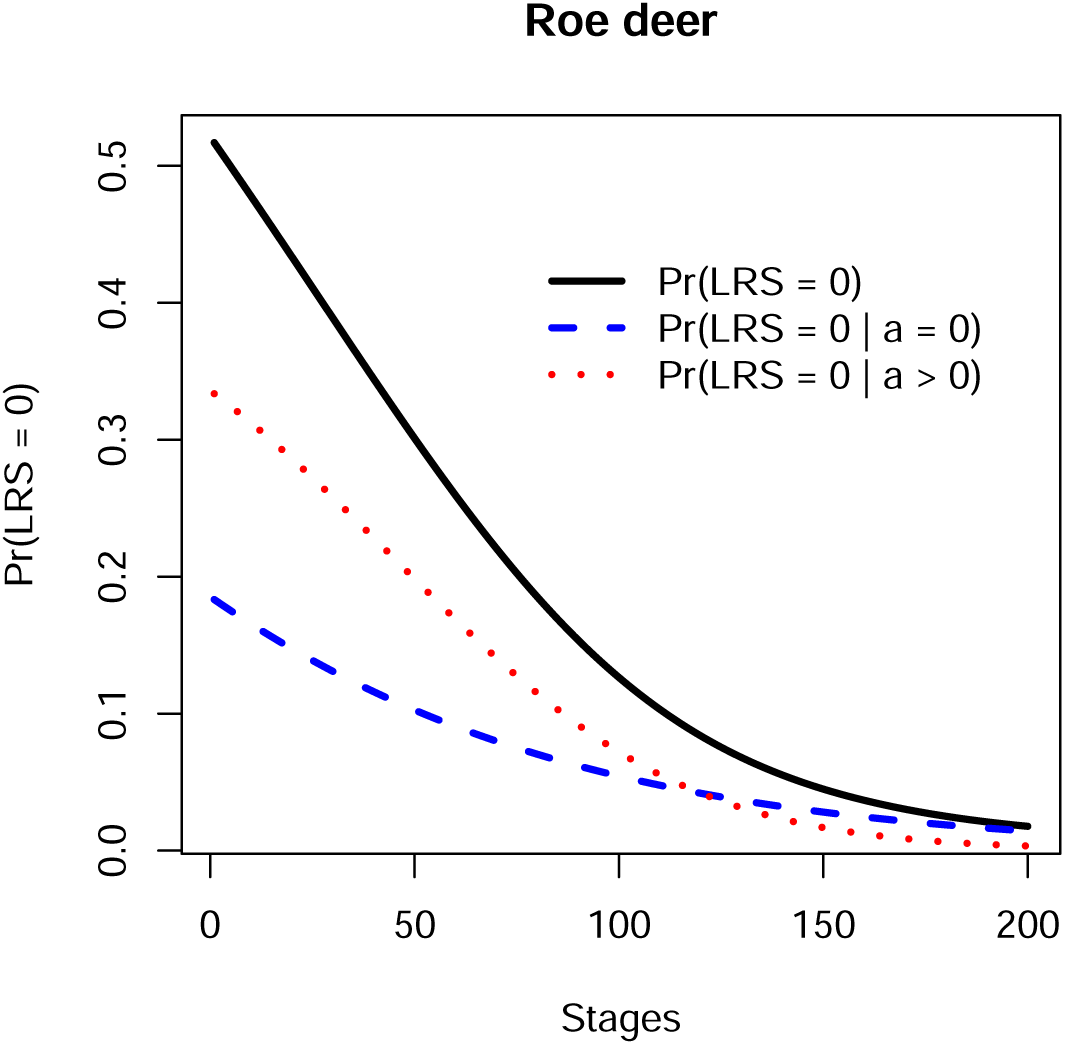
The probability that LRS = 0 for different initial size classes for yearlings of Roe deer *Capreolus capreolus*. The blue dashed line shows the yearling death probability. The red dotted line shows the probability of having no offspring if you live to later ages.

### A.11 The flow chart of decisions

**Figure A.4:**
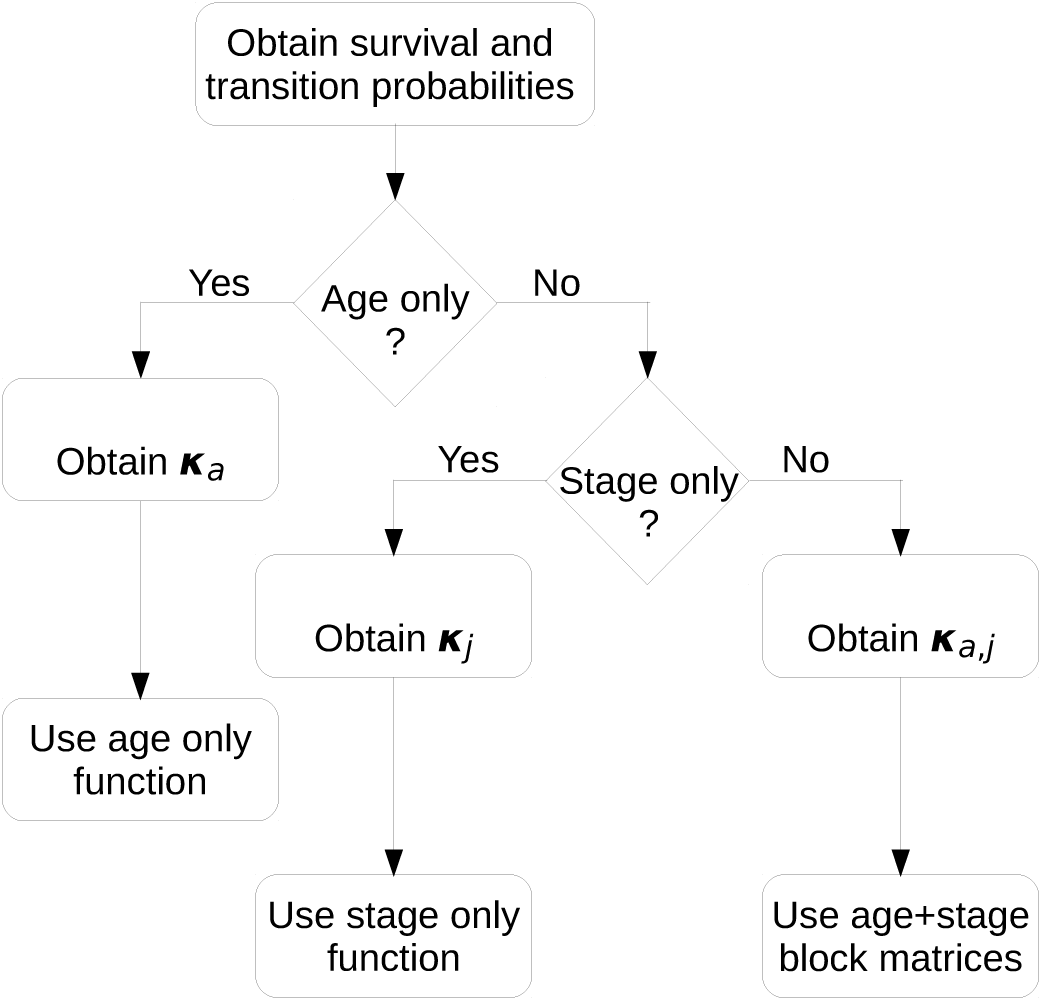
The flow chart of decisions. Survival probabilities (Table 1) and/or survival-transition matrices (Table 1); Reproduction distributions ***κ***_***a***_, ***κ***_***j***_, ***κ***_***a***,***j***_ (Table 1).

**Table A.1.**
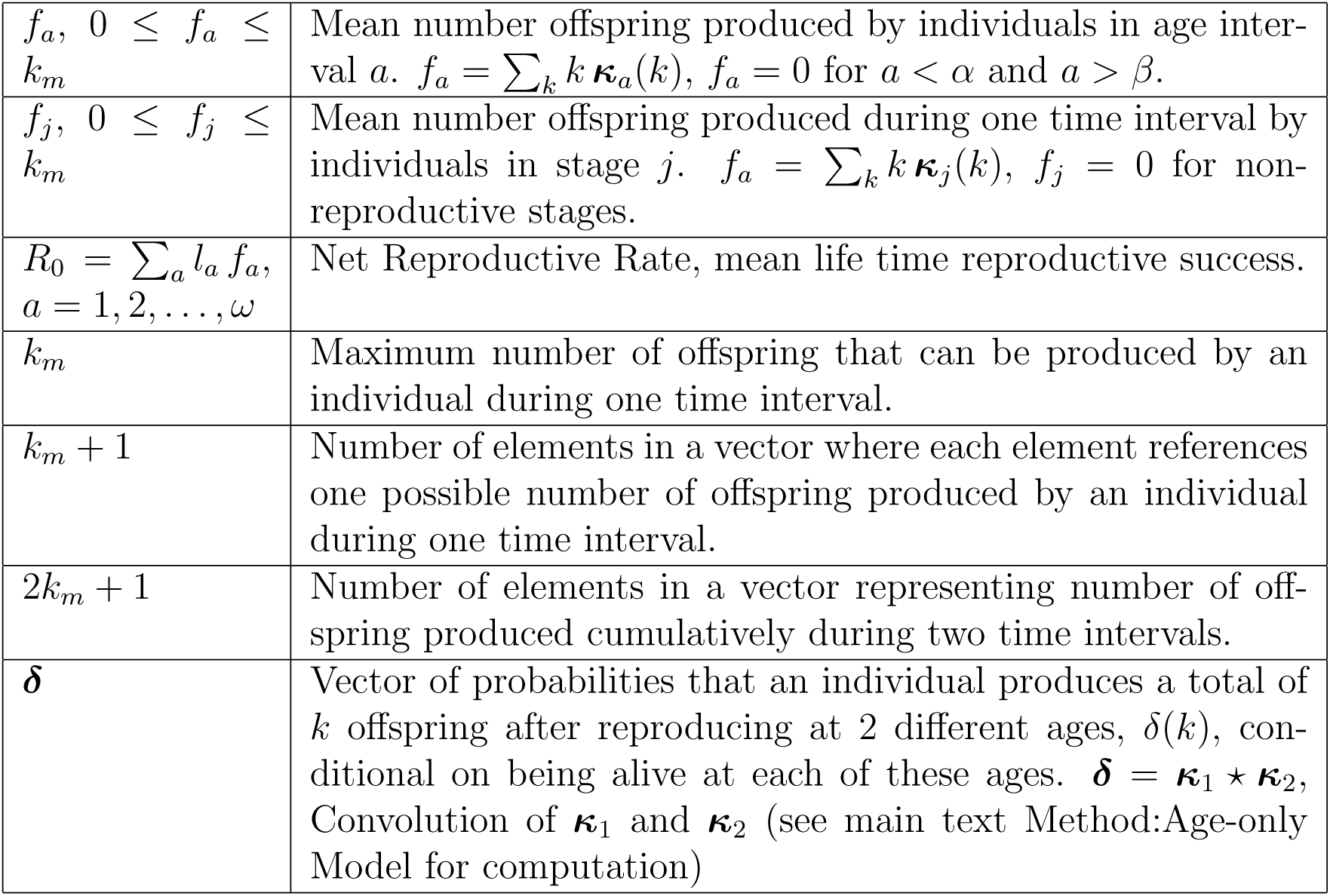
Definitions.

